# SC3s - efficient scaling of single cell consensus clustering to millions of cells

**DOI:** 10.1101/2021.05.20.445027

**Authors:** Fu Xiang Quah, Martin Hemberg

## Abstract

Technological advances have paved the way for single cell RNAseq (scRNAseq) datasets containing several million cells ^1^. Such large datasets require highly efficient algorithms to enable analyses at reasonable times and hardware requirements ^2^. A crucial step in single cell workflows is unsupervised clustering, which aims to delineate putative cell types or cell states based on transcriptional similarity ^3^. Here, we present a highly efficient k-means based approach, and we demonstrate that it scales linearly with the number of cells with regards to time and memory.

---

The most popular methods for unsupervised clustering of scRNAseq data are the Louvain and Leiden algorithms. They represent cells as a k-nearest neighbor graph where densely connected modules are identified as clusters ^4^. However, these methods can be biased by a poorly specified graph, running the risk of identifying structures that are not present in the data ^5^. More generally, as it can be shown that no single clustering algorithm will feature all desired statistical properties and perform well for all datasets, the field would benefit from having more methodologies available ^6^.

One of the most widely used unsupervised clustering in general is k-means clustering, and it forms the basis of the single cell consensus clustering (SC3) method ^7^. To achieve robust and accurate results SC3 uses a consensus approach whereby a large number of parameter combinations are evaluated and subsequently combined. However, both the k-means clustering and the consensus algorithm come at significant computational costs: both the run time and memory use scale more than quadratically with the number of cells, prohibiting application to large datasets.

Here, we present a new version of this algorithm, single cell consensus clustering with speed (SC3s), where several steps of the original workflow have been optimized to ensure that both run time and memory usage scale linearly with the number of cells (**Fig 1A, S1**). This is achieved by using a streaming approach for the k-means clustering ^8^ which makes it possible to only process a small subset of cells in each iteration. Each of the subsets can be efficiently processed at constant time and memory. In addition, as part of an intermediary step, which was not part of the original method, a large number of microclusters are calculated. The microclusters can be reused for different choices of *k*, and this allows substantial savings when analyzing multiple values of *k*, something that is very common in practice. We have also improved the consensus step by adopting a one-hot encoding approach ^9^, as opposed to the original co-association based method (**Fig S2**).

**Figure 1:**
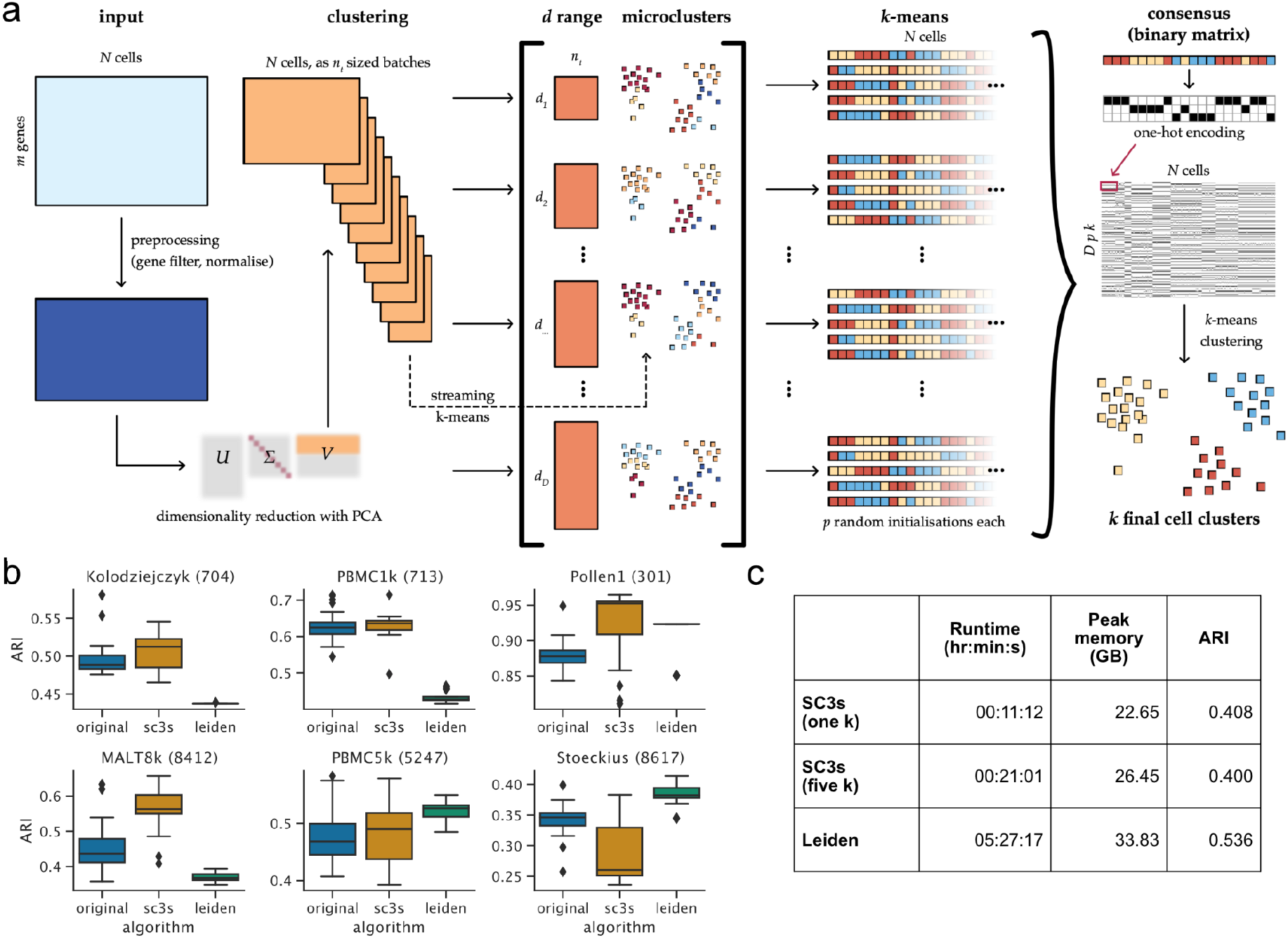
The SC3s framework for single cell consensus clustering. (a) Schematic depicting the overall flow of the algorithm. (b) Adjusted Rand index (ARI) accuracy benchmarks on gold-standard datasets with <10,000 cells. Boxplots show the distribution across 25 realisations of each algorithm. Numbers in parentheses denote the cell count in the specified dataset. (c) Runtime, memory and ARI performance benchmarked on the 2 million mouse organogenesis cell atlas dataset (average of 5 runs).

To evaluate the accuracy of SC3s we used six datasets with <10,000 cells where the cell labels have been defined using orthogonal methods, allowing us to compare the results of the transcriptome clustering to a ground truth ^7^ (**Table S1**). These benchmarks show that SC3s has an accuracy which is comparable to the original algorithm (**Fig 1B**). Furthermore, the performance is robust across a broad range of user-customisable parameters (**Fig S3–S5**). Finally, SC3s compares favourably against the Leiden algorithm under different parameterisations in terms of accuracy and runtime (**Fig 1B, S1, S6**).

To examine the performance for large datasets, SC3s was benchmarked on the mouse organogenesis cell atlas dataset which contains 2,026,641 cells ^1^. Processing, filtering and dimensionality reduction were performed as in the original publication, after which the clustering performance of SC3s was compared against the Leiden algorithm. Although the peak memory usage was comparable, SC3s was ~18 times faster (20 mins vs 6 hrs), even when evaluating five *k* values (**Fig 1C**). The slightly lower accuracy was expected because cell labels used for comparison originated from the Louvain algorithm, a method very similar to the Leiden algorithm, making them an imperfect ground truth. Visual inspection of the assigned labels also revealed that SC3s was able to capture the major structures identified by the authors (**Fig S7**).

Overall, SC3s is a major improvement over its predecessor, and it represents a scalable and accurate alternative to the widely used neighbourhood graph clustering methodologies. Moreover, it is integrated with the popular Scanpy package ^10^, making it easy to incorporate into existing workflows. Thus, SC3s will allow researchers to analyze scRNAseq datasets as they scale to millions of cells.

## Data availability

All datasets used for benchmarking are available publically, and they are listed in Supplementary Table S1.

## Code availability

The Python code for SC3s is licensed under a BSD-3 Clause License. Instructions to install from pip and conda channels are available on GitHub: https://github.com/sc3s/sc3s.

## Benchmarking setup

Benchmarks were run on OpenStack virtual machines with four Intel Core Processor Broadwell vCPUs and 64GB RAM.

## Acknowledgements

FXQ was supported by a Wellcome Trust PhD studentship. MH was funded by a core grant from the Wellcome Trust. We would like to thank the Cellular Genetics Informatics team at the Wellcome Trust Sanger Institute for providing compute resources, particularly Simon Murray for helping package SC3s.

## Author contributions

The project was conceived by FXQ and MH. FXQ wrote the code and analysed the data. MH supervised the research. FXQ and MH wrote the manuscript.

---

**Figure S1:**
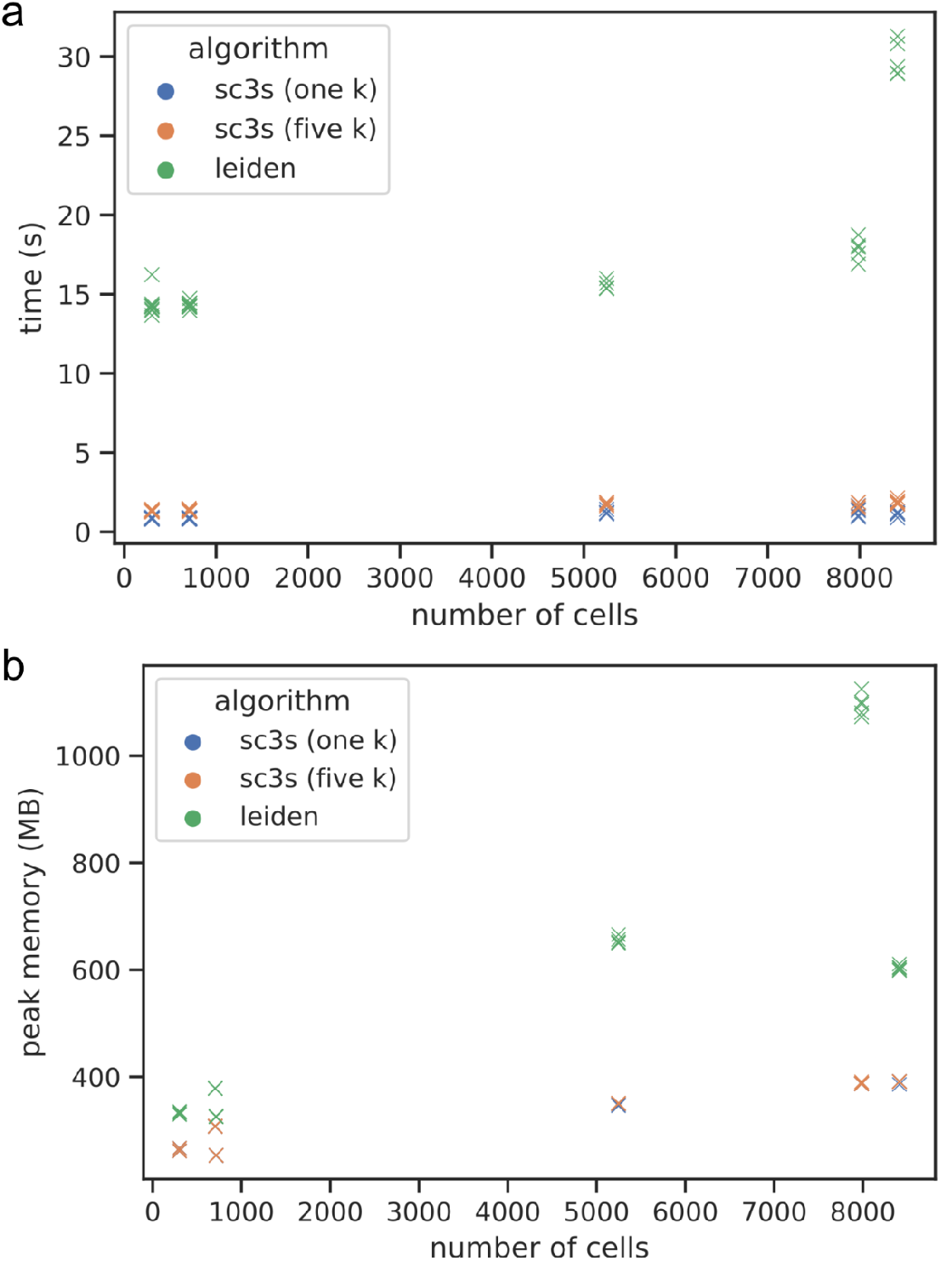
Runtime and memory performance relative to the number of cells. For each dataset, five realisations of the SC3s (with one and five *k* values tested) and Leiden (build graph + cluster cell steps) algorithms were plotted.

**Figure S2:**
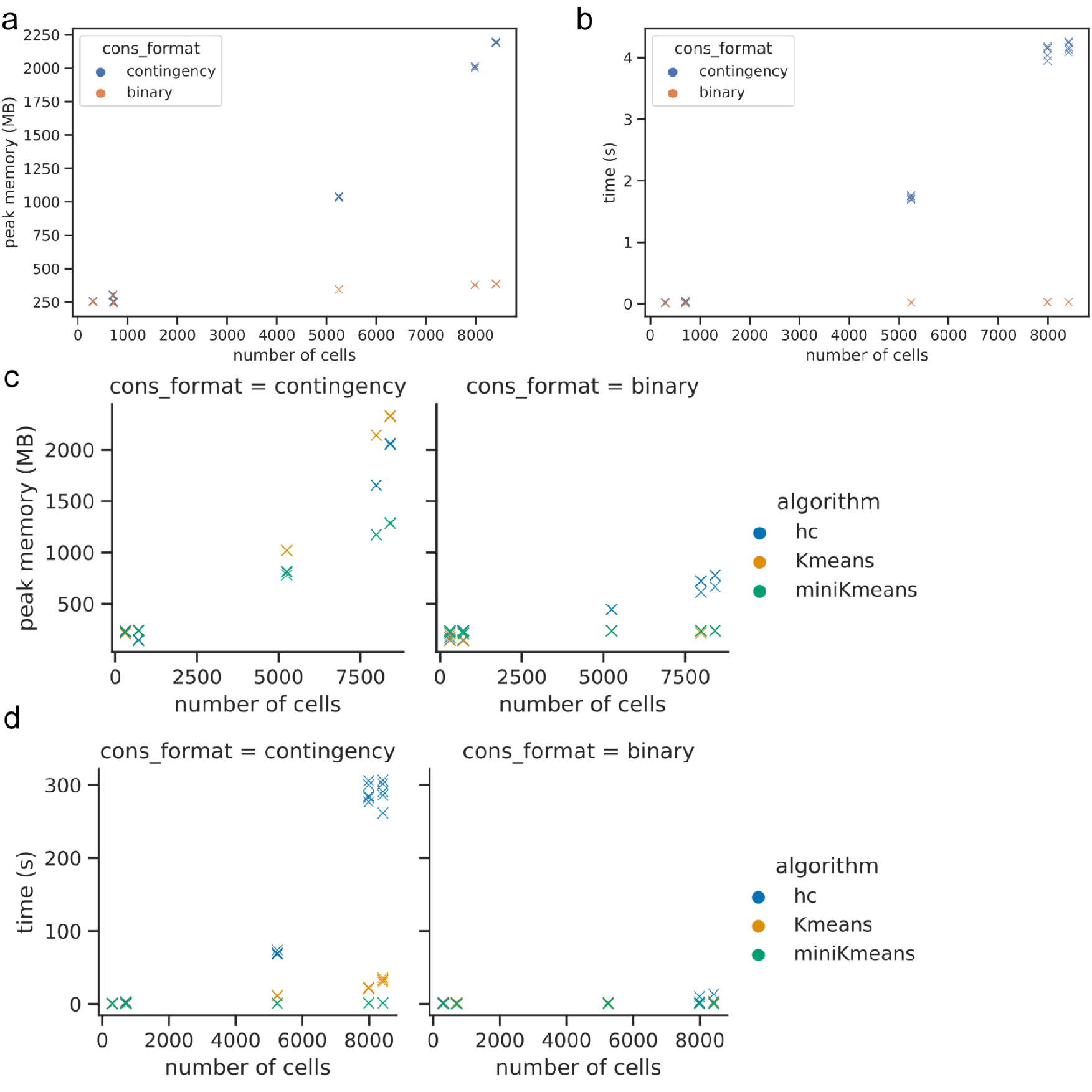
Performance improvements of the one hot encoding approach for consensus clustering. Runtime and memory usage is significantly reduced not only for the construction of the consensus matrix (a, b), but also its subsequent use to generate the final clusters (c, d).

**Figure S3:**
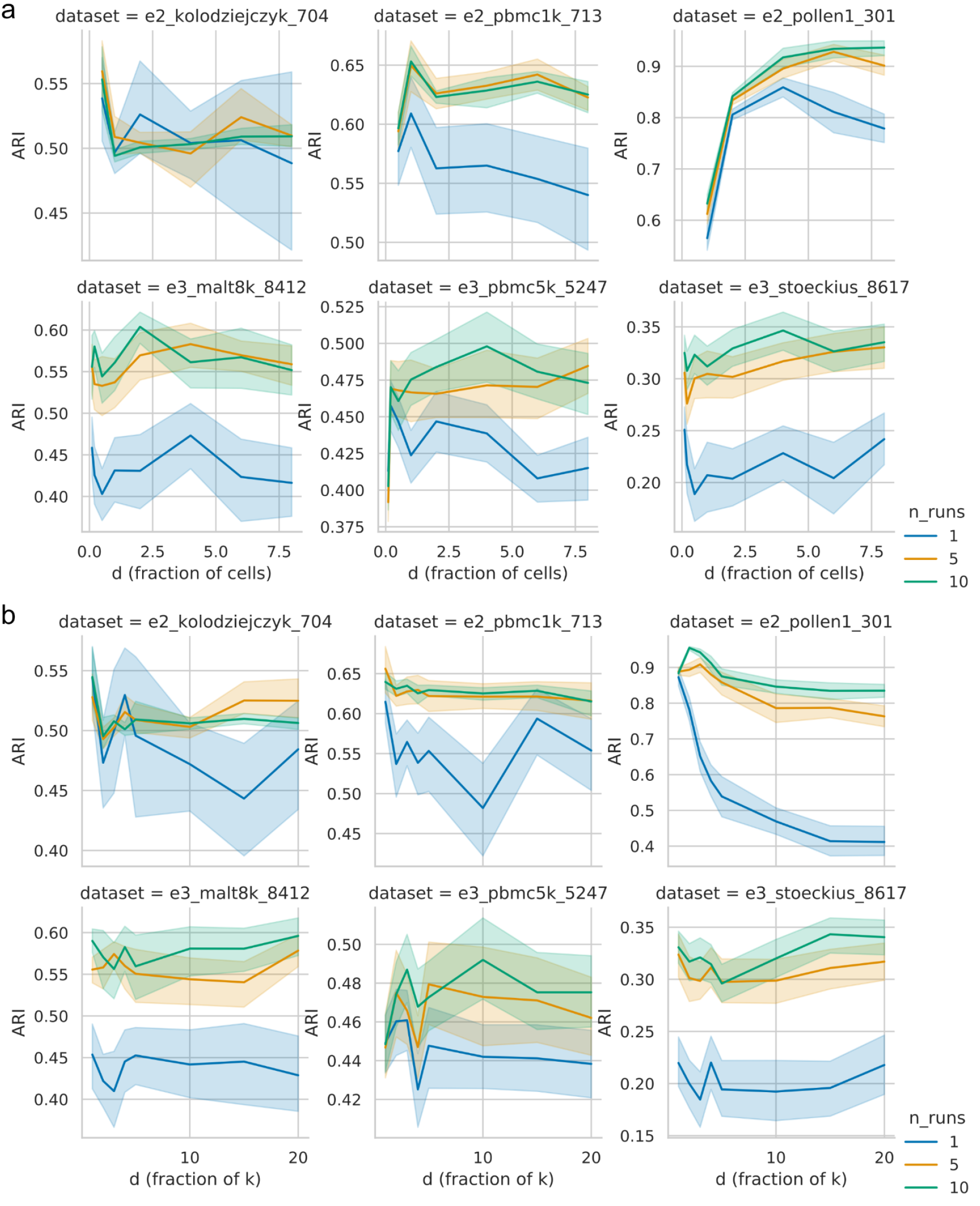
Stability of SC3s clustering results across a wide range of d values (number of PCs). The mean ARI performance with 95% confidence intervals is shown across 25 independent realisations for each *d* value. *d* values are plotted as a fraction of the true number of clusters k (panel a), and as a fraction of the number of cells (panel b). In general, a plateau of robust ARI performance is obtained once a sufficient number of PCs are included. The algorithm exhibits higher accuracy and more stability when more runs are used to obtain the consensus.

**Figure S4:**
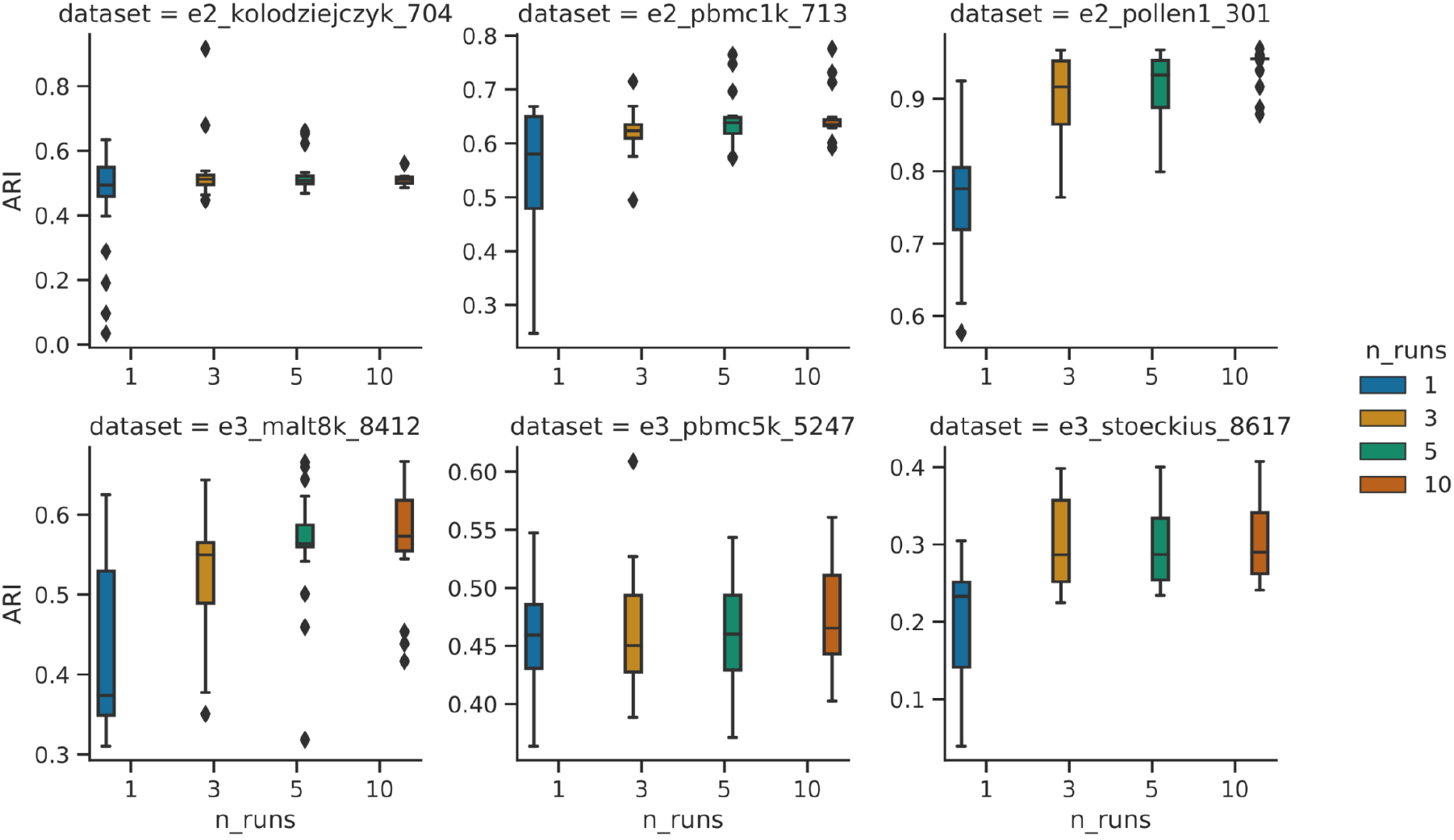
Stability of SC3s clustering results with respect to the n_runs parameter. *n_runs* denote the number of runs used to calculate the consensus. ARI performance of SC3s increases with a higher number of iterations run to obtain the consensus. 25 independent realisations were used for each dataset.

**Figure S5:**
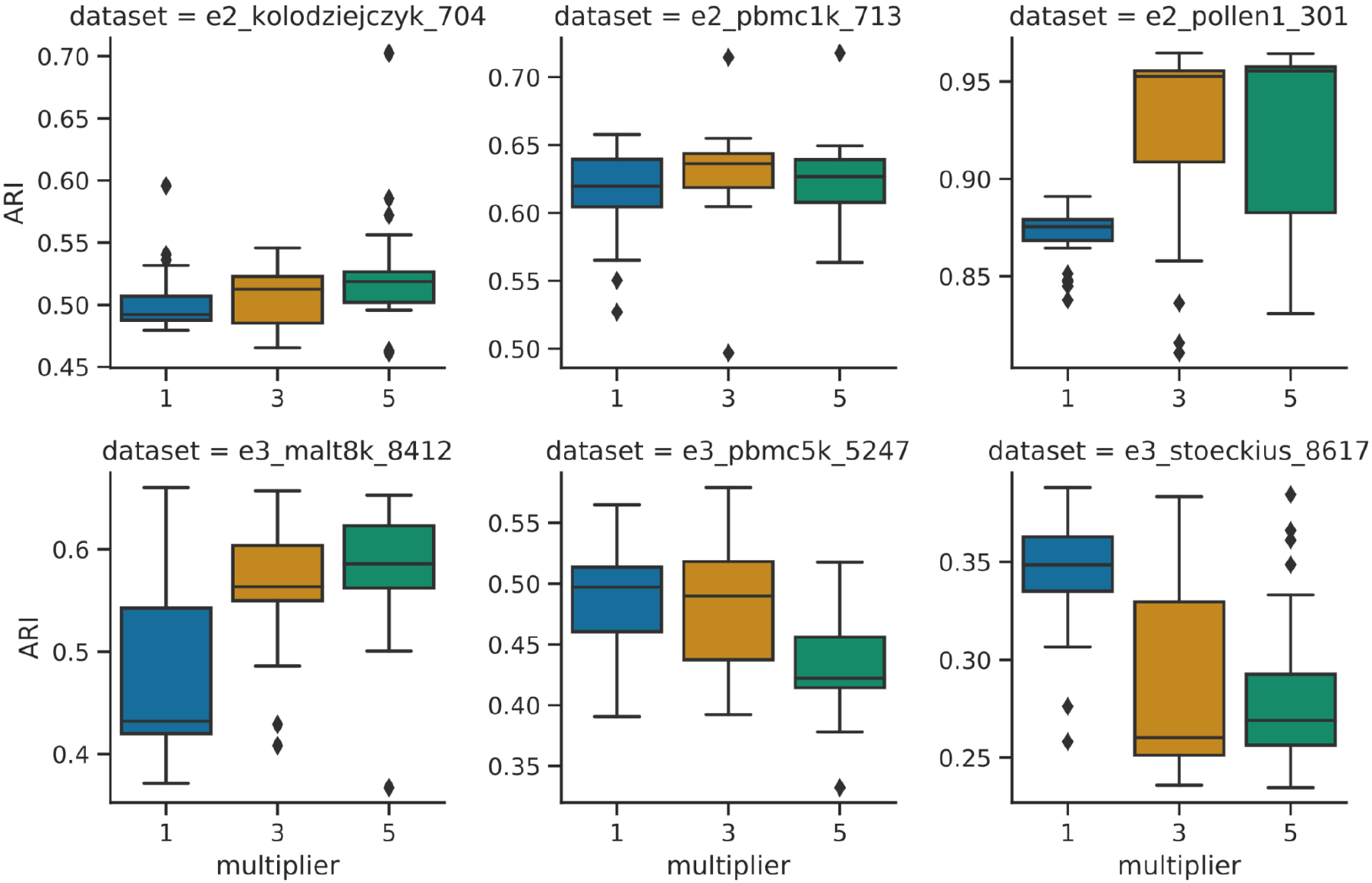
Stability of SC3s clustering results with respect to the multiplier parameter. The ARI performance is examined under a different number of microclusters, which was calculated as the true number of *k* multiplied by the multiplier value. 25 independent realisations were used for each dataset.

**Figure S6:**
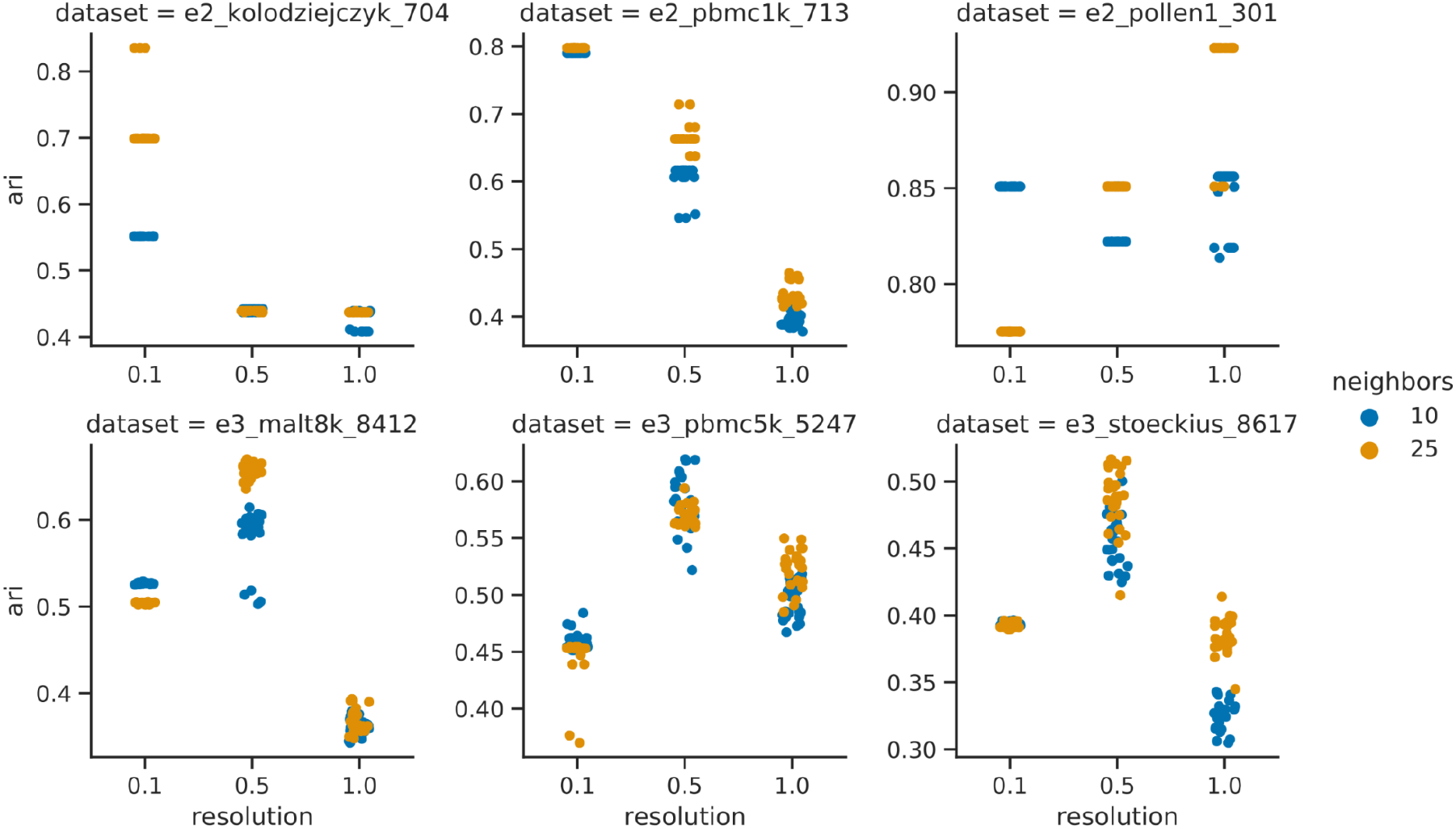
ARI performance of Leiden algorithm, under different parameterisations. Color denotes the number of neighbours used to construct the neighbourhood graph, while the x-axis denotes the resolution parameter in the Leiden algorithm. 25 independent realisations each.

**Figure S7:**
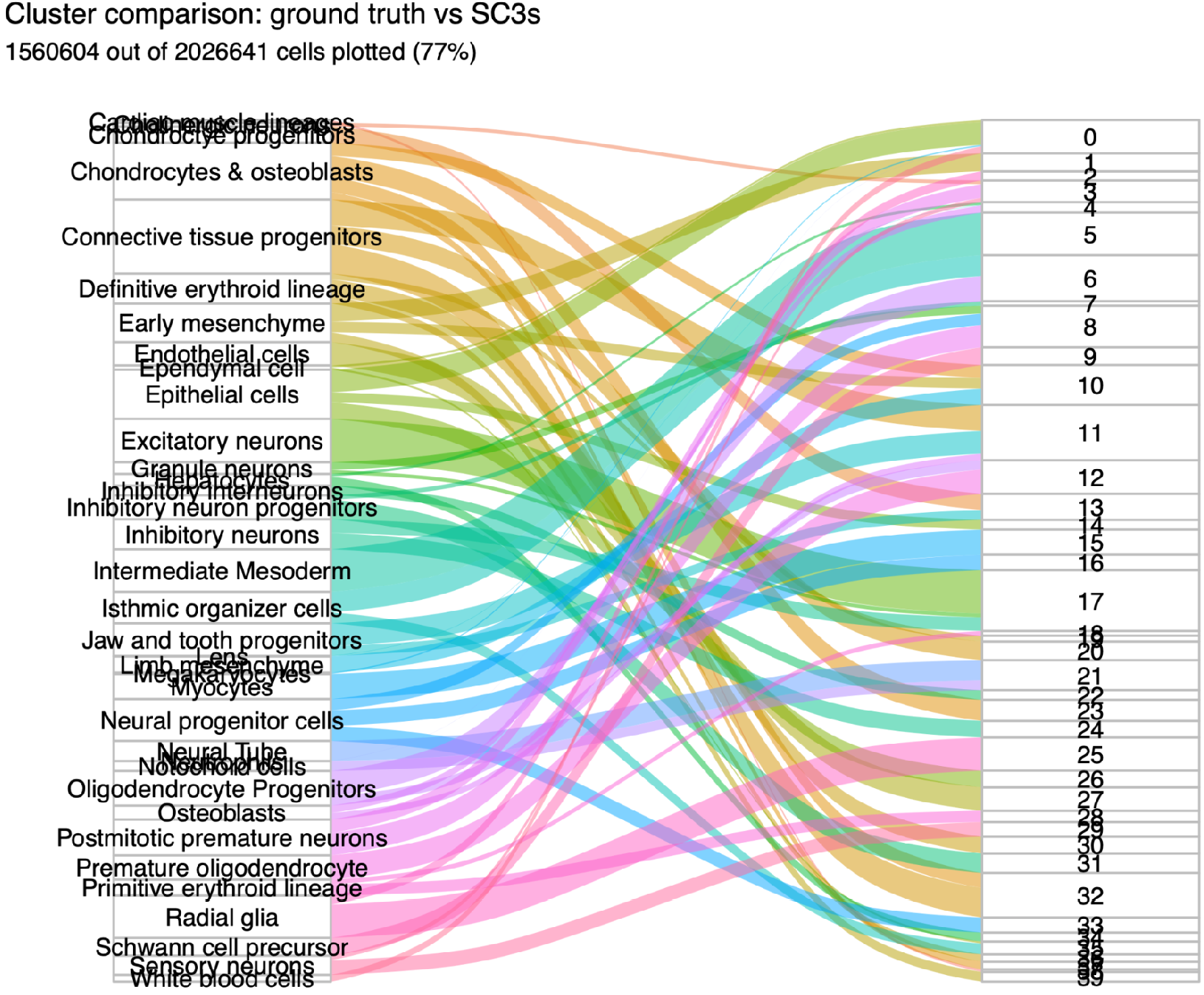
Sankey diagram comparing the clusters reported by Cao et al (left) with clusters assigned by SC3s (right). The width of the edges correspond to the number of cells common to the linked nodes. To focus only on the major structure in the data, only edges which contain above 20% of the cell types in one of the nodes were included.

**Table S1:** Datasets used for benchmarking.

| Name | Identifier | Cell count | True k | Source | URL |
| --- | --- | --- | --- | --- | --- |
| Kolodziejczyk | e2_kolodziejczyk_704 | 704 | 3 | Kolodziejczyk, A. A. *et al.* <sup>1</sup> | https://hemberg-lab.github.io/scRNA.seq.datasets/mouse/esc/#kolodziejczyk |
| PBMC1k | e2_pbmc1k_713 | 713 | 6 | 10x Genomics | https://support.10xgenomics.com/single-cell-gene-expression/datasets/3.0.0/pbmc_1k_protein_v3 |
| Pollen | e2_pollen_1 | 301 | 11 | Pollen A.A. *et al.* <sup>2</sup> | https://hemberg-lab.github.io/scRNA.seq.datasets/human/tissues/#pollen |
| MALT8k | e3_malt8k_8412 | 8412 | 8 | 10x Genomics | https://support.10xgenomics.com/single-cell-gene-expression/datasets/3.0.0/malt_10k_protein_v3 |
| PBMC5k | e3_pbmc5k_5247 | 5247 | 8 | 10x Genomics | https://support.10xgenomics.com/single-cell-gene-expression/datasets/3.0.2/5k_pbmc_protein_v3 |
| Stoeckius | e3_stoeckius_8617 | 8617 | 8 | Stoeckius, M. *et al.* <sup>3</sup> | https://www.ncbi.nlm.nih.gov/geo/query/acc.cgi?acc=GSE100866 |
| MOCA | e6_moca_2mil | 2026641 | 38 | Cao, J. *et al.* <sup>4</sup> | http://oncoscape.v3.sttrcancer.org/atlas.gs.washington.edu.mouse.rna |

1. Kolodziejczyk, A. A. *et al.* Single Cell RNA-Sequencing of Pluripotent States Unlocks Modular Transcriptional Variation. *Cell Stem Cell* **17**, 471–485 (2015). 2. Pollen, A. A. *et al.* Low-coverage single-cell mRNA sequencing reveals cellular heterogeneity and activated signaling pathways in developing cerebral cortex. *Nat. Biotechnol.* **32**, 1053–1058 (2014). 3. Stoeckius, M. *et al.* Simultaneous epitope and transcriptome measurement in single cells. *Nat. Methods* **14**, 865–868 (2017). 4. Cao, J. *et al.* The single-cell transcriptional landscape of mammalian organogenesis. *Nature* **566**, 496–502 (2019).

